# Leukemic fusion genes repress viral gene expression and expel adenovirus from persistently infected human B lymphocytes but evidence of the virus lingers behind

**DOI:** 10.64898/2025.12.19.695471

**Authors:** H. T. Wilms, D. A. Ornelles, L. R. Gooding, N. Maganti, C. Garnett-Benson

**Affiliations:** Department of Biology, Georgia State University, Atlanta, GA 30303; Department of Microbiology and Immunology, Wake Forest School of Medicine, Winston-Salem, NC 27157; Emory University School of Medicine, Department of Microbiology and Immunology, Atlanta, Georgia 30322

**Keywords:** leukemia, adenovirus, species C, viral persistence, human lymphocytes, hit and run

## Abstract

Species C adenoviruses infect virtually all children in the first few years of life. These viruses can establish asymptomatic persistent infections in mucosal-associated lymphocytes and can infect lymphocytes *in utero*. Although adenovirus is a DNA tumor virus, it has not yet been shown to initiate cancer in humans. Epidemiological studies of B cell precursor acute lymphoblastic leukemia (ALL), point towards an infectious etiology. The ETV6/RUNX1 fusion protein results from a chromosomal translocation t(12; 21) believed to initiate ALL. This translocation can be detected *in utero* and has been identified in as much as 40% of clustered cases of leukemia, which are those most likely to have been initiated by an infectious agent. Infectious agents have not yet been detected in leukemic cells, causing speculation that the oncogenic agent has been lost in the transformed progeny through a “hit and run” mechanism. In the current study, we attempt to model the “run” of adenovirus in B-lymphocytes by forcing expression of leukemic fusion genes to determine if they create an environment that is refractory to adenovirus persistence. Here we show that the common leukemic fusion proteins, ETV6/RUNX1 or RUNX1/MTG8, reduce adenovirus persistence in a B-lymphocyte line but do not dampen the initial acute infection phase. Further, we show that ETV6/RUNX1 can bind to the viral genome and that some viral gene expression appears to be suppressed through the activities of HDACs in ETV6/RUNX1-expressing cells. Finally, we show that the expression of virally silenced cellular genes remains repressed even after the loss of the virus from infected cells. The results of the current study provide support for how adenovirus could be lost from translocation containing lymphocytes and provide evidence that adenovirus can leave a lasting imprint on cells previously infected in the form of an epigenetic echo.

## Introduction

Adenoviruses are ubiquitous non-enveloped, icosahedral viruses (1). The dozens of serotypes that infect humans are assigned to seven species, the most common of which include species C serotypes 1, 2, 5 and 6. These viruses establish lytic infections in epithelial cells of the upper respiratory and gastrointestinal tracts (1) and latent infections in T and B lymphocytic cells (2, 3). The peak incidence of species C viruses in human mucosal lymphocytes occurs between two and five years of age (2, 4). Adenovirus infections also occur *in utero* and the species C viruses have been detected in the cord blood of one in 25 newborns (5).

Primary rodent cells are readily immortalized by the oncogenes of adenovirus (6, 7). Human mesenchymal stromal cells have been shown to be susceptible to transformation by the adenovirus E1 oncogenes (8). Although adenovirus is not known to cause any human cancer, an oncogenic role for this small DNA tumor virus is not unreasonable. Adenoviral gene products inactivate critical cell cycle checkpoints such as p53 and pRb, whose loss is known to contribute to tumor progression (7, 9–14). Adenovirus suppresses cellular DNA-break repair pathways in order to preserve the integrity of the linear double-stranded viral genome (15, 16). The loss of DNA-break repair promotes genomic instability (15) which is another hallmark of cancer (14). The virus has also been shown to epigenetically repress the expression of several cellular genes with proposed tumor suppressor gene capacity (17). Additionally, adenovirus elicits re-replication of cellular DNA in quiescent cells, which can lead to hyperdiploidy (18). Thus, adenovirus could contribute to cancer development either by disrupting DNA repair processes or by repression of tumor suppressor genes that help promote transformation. Despite its oncogenic potential, adenovirus has yet to be implicated as the cause of any human cancer. One hypothesis for the absence of any connection between adenovirus and human cancers is that oncogenic mutations elicited by the virus preclude retention of the virus in the affected cell. This is the premise behind the “hit-and-run” model of viral oncogenesis (19, 20). The loss of adenoviral genes (19) and the loss of herpes simplex virus gene products from rodent cells transformed *in vitro* (21) by these viruses is experimental evidence that supports this possible mechanism of cellular transformation.

Viral genes are expressed in human cancers caused by Epstein-Barr virus (EBV) and human papillomavirus (HPV) (22, 23). Cancers caused by these viruses emerge from tissue that harbors a persistent or latent infection. EBV is responsible for tumors of B-cell origin including Hodgkin’s lymphoma and Burkitt’s lymphoma and tumors of the epithelial origin including nasopharyngeal carcinoma and gastric cancer (22). All of these tumors are latently infected with EBV and express a subset of the viral genes (22, 24). HPV may establish quiescent, clinically latent infections in long-lived epithelial cell progenitors (25). Virtually all oropharyngeal cancers, as well as cancers of the uterine cervix, that are caused by HPV express the viral HPV E6 and E7 oncogenes (26, 27). Because adenovirus can chronically infect B lymphocytes, we and others have speculated that this virus could initiate events that lead to the development of childhood acute lymphoblastic leukemia (ALL) (28, 29) by a mechanism that results in the loss of the viral genome from the transformed cell since no viral products have been found associated with the cancerous cells.

Epidemiological evidence supports an infectious contribution to development of childhood ALL. In particular, common precursor B cell ALL has been under intense investigation for an infectious cause due to the marked pattern of disease clustering (30–32). Because no infectious agent has yet been identified within the leukemic cells themselves (33), the viral contribution to transformation may be occurring through a hit-and-run mechanism. ALLs are typically associated with identifiable chromosomal translocation mutations that differ between childhood and adult leukemias (34, 35). Indeed, the best evidence for a multistep process leading to development of childhood ALL comes from the detection of leukemia-associated translocations in the blood of newborns many years before the development of overt leukemia (36, 37). The most common translocation in childhood precursor B cell ALLs, t(12; 21) which produces the fusion protein ETV6/RUNX1, is found in about one quarter of cases (35). Moreover, a recent study found that when only “clustered” cases of leukemia are considered (those most likely to have been initiated by an infectious agent) the frequency of the *ETV6/RUNX1* translocation rises to 40% (38).

While mutations in *RUNX1* frequently contribute to the initiation of leukemia (39, 40), RUNX1 it is an important transcription factor in normal B lymphocyte development (39–43). The RUNX1 proteins show a distinct relationship to adenovirus. Human RUNX1a and RUNX1b isotype variants allowed nuclear localization of the normally cytoplasmic adenovirus E1B-55K protein in a mouse cell line where the RUNX1 proteins localized to sites enriched for viral RNA processing factors. This association displaced the adenovirus E4orf6 protein from the complex. These results suggest that RUNX1 may play a role in adenoviral RNA processing (44). RUNX1, and RUNX1 leukemic fusion proteins, are also known to bind to DNA and alter gene transcription (39). Interestingly, human species C adenoviruses have canonical RUNX1-binding motifs in the promoter regions of early genes that control viral retention and replication (45–47). Wild-type RUNX1 is reported to both activate and repress cellular gene expression by recruiting histone acetyl transferases (HATs) or histone deacetylases (HDACs), respectively. By contrast, the ETV6/RUNX1 fusion protein primarily represses gene expression by HDAC recruitment (39, 48). Little is known about the influence of RUNX1, or RUNX1 containing fusion proteins, on persistent or latent infections in B cells. We have previously shown that B cell lines naturally harboring the *ETV6/RUNX1* chromosomal translocation had reduced infectivity with species C adenovirus (17). Thus, we reasoned that the cells containing this translocation could be refractory to persistent infections with the virus.

If adenovirus contributes to childhood precursor-B cell leukemia, the absence of any viral product in these cancers would imply the existence of a hit-and-run mechanism. However, no study has yet shown a mechanism for viral loss from precancerous cells. Because of the difficulty of measuring the loss of a virus from the associated diseased tissue in humans, we sought to model this behavior experimentally. We asked if the transcriptionally repressive leukemic fusion proteins (39, 48) could inhibit adenovirus retention in persistently infected B cells. Our results show for the first time that both viral gene expression and the viral genome are lost during a persistent infection of B cells expressing *RUNX1* leukemic fusion genes. Loss of expression of the viral DNA binding protein precedes the loss of the viral genome from ETV6/RUNX1 expressing cells. The ETV6/RUNX1 fusion protein binds to viral DNA and reduced expression of some viral genes appears to be mediated by histone deacetylation. Moreover, we found that expression of the cellular genes *BNIP3*, *CXADR*, and *SPARCL1,* which are epigenetically repressed in persistently infected cells, (17), remains depressed even after loss of the virus from the cells. Our finding reveal that the *RUNX1* leukemic fusion products may expel adenovirus from persistently infected B cells while an epigenetic echo of the infection remains in the cells.

## Materials and methods

### Cell lines

The human cell A549 lung carcinoma cell line was purchased from the American Type Culture Collection (ATCC, Manassas, VA). BJAB (EBV-negative Burkitt’s lymphoma) cells were obtained from the ATCC (49). BJAB cells were grown in RPMI medium supplemented with 10% fetal calf serum (FCS) and 10 mM glutamine. A549 cells were grown in Dulbecco’s modified Eagle medium (DMEM) with 4.5 μg of glucose per ml, 10% FCS, and 10 mM glutamine. Cells were tested by Genetica for cell line authentication and were routinely tested to ensure the absence of mycoplasma.

### Creation of stable cell lines

The original expression plasmids for the *ETV6/RUNX1* and *RUNX1/MTG8* were kindly provided by S. Hiebert (Vanderbilt University) and were subcloned into pTARGET (Promega) using the *Eco*RI restriction enzyme. A549 cells were transfected using Lipofectamine LTX with Plus reagent (ThermoFisher Scientific). BJAB cells were transfected using electroporation as described in Mchichi *et at* (50), with minor modifications. Briefly, 4x10^6^ cells in cold serum-free RPMI were transfected with 2μg of plasmid by electroporation at 120 V, 960 μF using cuvettes with an inner width of 2 mm. After 20 min of recovery at 37°C, cells were plated into 6-well plates in RPMI supplemented with 10% FBS at 37°C for 48 hrs. Cell cultures were then supplemented with 1 mg per ml of G-418 (ThermoFisher Scientific) for at least 21 days before use in experiments. G-418 was maintained at 0.5 mg per ml throughout experiments to maintain stably transfected cell lines.

### Adenoviruses

Wild-type species C Ad5 adenovirus was obtained from William S. Wold (St. Louis University). Similarly, the phenotypically wild-type mutant virus, Ad5dl309, was obtained from Tom Shenk (Princeton University, Princeton, NJ. Ad5dl309 is an Ad5 mutant that lacks the genes for the E3 RIDα and RIDβ proteins as well as the 14,700-molecular-weight protein (14.7K protein) (51).

### Infection of lymphocytes with adenovirus

Infection of lymphocyte cell lines with adenovirus was performed as described previously (52) with minor modifications. Lymphocytes were collected and washed in serum-free (SF) RPMI medium. Cell density was adjusted to 10^7^ cells per ml in SF-RPMI medium. Virus was added to the cell suspension at 50 PFU/cell, spun for 4 5min at 1000 x g at 25°C, resuspended by agitation. Cells were then incubated at 37°C for 1.5 hrs with gently flicking every 30 min. The infected cells were washed three times with RPMI complete medium and then resuspended in RPMI complete medium at 5x10^5^ cells per ml. Cell concentration and viability were monitored throughout the infection.

### Reverse transcription and quantitative PCR analysis of viral and cellular mRNA levels

RT-qPCR was performed as described previously (17), with minor modifications. Briefly, total RNA was isolated from cells using the RNeasy Mini Kit (Qiagen Inc. Valencia, CA). RNA was treated with Rnase-free DNase (Qiagen) on isolation columns and quantified. Greater than 100 ng were reverse transcribed (RT) into cDNA, in 20 μL reaction volumes, using Maxima First Strand cDNA Synthesis Kit (ThermoScientific). RT-enzyme negative controls were included for each reaction. Primers and probes were obtained from Integrated DNA Technologies (Coralville, IA). EIF1 and viral primer and probe sequences used are below. Primer and probes for the cellular genes *CXADR, BNIP3,* and *SPARCL1* were used as previously described (17). Probes were labeled at the 5’ end with 6-carboxyfluorescein (FAM) reporter molecule and typically contained dual ZEN and Iowa Black quenchers. Each sample was run in duplicate with at least 2 experimental repeats for each virus tested. All analyses were performed via the comparative threshold cycle (Ct) method after 40 cycles (53). Target Cts were normalized to the EIF1 housekeeping gene. In some experiments, HDAC inhibition was performed by treating cells with 175 nM Trichostatin A (TSA) followed by quantifying transcript levels (17).

E1A (Sense sequence, 5’- GTTAGATTATGTGGAGCASCCC-3’, anti-sense sequence, 5’-CAGGCTCAGGTTCAGACAC -3’, probe sequence, 5’-6 FAM-ATGAGGACCTGTGGCATGTTTGTCT-3IABkFQ-3’)

E2A (Sense sequence, 5’-GAAAACTTCACCGAGCTGC-3’, anti-sense sequence, 5’-ACACGTTGCGATACTGGTG-3’, probe sequence, 5’-6 FAM-CGGATGGTTGTGCCTGAGTTTAAGTG-3IABkFQ-3’)

E3GP19K (Sense sequence, 5’-TTTACTCACCCTTGCGTCAG-3’, anti-sense sequence, 5’-GCAGCTTTTCATGTTCTGTGG-3’, probe sequence, 5’-6 FAM-CTGGCTCCTTAAAATCCACCTTTTGGG-3IABkFQ-3’)

TLP HEXON (Sense sequence, 5’-AAAGGCGTCTAACCAGTCAC-3’, anti-sense sequence, 5’-CCCGAGATGTGCATGTAAGAC-3’, probe sequence, 5’-6 FAM-CGCTTTCCAAGATGGCTACCCCT-3IABkFQ-3’)

EIF1 (Sense sequence, 5’- GATATAATCCTCAGTGCCAGCA-3’, anti-sense sequence, 5’-GTATCGTATGTCCGCTATCCAG-3’, probe sequence, 5’-6 FAM-CTCCACTCTTTCGACCCCTTTGCT-3IABkFQ-3’)

### Quantitative real time PCR analysis of viral DNA levels

Infected or uninfected control cells were washed in phosphate-buffered saline (PBS) and lysed in 100 μl of NP-40–Tween buffer containing proteinase K, as described in Garnett et al (2). Samples were tested by real-time PCR for a region of hexon gene that is conserved among species C adenovirus serotypes and the endogenous cellular gene glyceraldehyde-3-phosphate dehydrogenase (GAPDH). Samples were run in duplicate for each independent experiment. Data was analyzed using the comparative threshold cycle (Ct) method in some experiments, as described above. The amount of *hexon* gene DNA was normalized to the amount of GAPDH and then relative amounts were compared to the time point representing the peak of the acute infection. In some experiments, viral genome numbers were quantified using a standard curve, as previously described (2).

### RT-PCR detection of ETV6/RUNX1 and RUNX1/MTG8 transcripts

Total RNA was isolated from cells transfected with plasmids expressing either the *ETV6/RUNX1* or *RUNX1/MTG8* fusion genes and converted into cDNA as described above. The presence of ETV6/RUNX1 and RUNX1/MTG8 mRNA transcripts was tested by PCR amplification using indicated primers and PCR conditions from van Dongen *et al* and amplicons were run on a 2% agarose gel (54).

### Flow cytometry

Intracellular staining for the viral capsid protein, hexon, was used to detect productively infected cells by flow cytometry as previously described (55). Briefly, a mouse monoclonal antibody to adenovirus hexon protein (IgG1 κ, MAB8051, Chemicon International/Millipore) was used as a primary antibody. A mouse isotype IgG1, κ, antibody was used as a negative control for primary antibody staining (BD Pharmingen). Cells were subsequently stained with a goat anti-mouse IgG-APC conjugated secondary antibody (Life Technologies). Results were analyzed on a LSR Fortessa flow cytometer using FACSDiva Software (BD Biosciences). Isotype control staining was used to define the hexon-positive staining cells and was 5% or less for all samples evaluated.

### Immunoblots for protein detection

A total of 4-6 x 10^6^ cells were collected and washed in cold PBS, the cell pellet was resuspended in 1 ml of cold RIPA buffer (R0278, Sigma) supplemented with 1 mM EDTA (161-0729, BioRad), protease/phosphatase inhibitor (1861281, Thermo Scientific), and incubated on ice for 30 min. Samples were then sonicated briefly, and boiled for 5 min with equal amounts of 2X Laemmli sample buffer (161-0737, Bio-Rad) before being run on a SDS-PAGE gel. The separated proteins were transferred to a nitrocellulose membrane (BioRad). Immunoblotting of RUNX1 and RUNX1 fusion proteins was performed by using primary polyclonal rabbit anti-RUNX1 antibodies (ab23980, Abcam or PC285, CalBiochem) and secondary goat anti-rabbit IgG-HRP antibodies (sc-2004, Santa Cruz Biotechnology). Primary mouse antibody to actin (MAB1501, Chemicon) and secondary donkey anti-mouse IgG-HRP (sc-2314, Santa Cruz Biotechnology) was blotted in parallel as a protein loading control Proteins were visualized with HyGLO Reagent A/B (E2500, Denville Scientific Inc) and visualized with HyBlot ES Autoradiography Film (E3218, Denville Scientific Inc). The adenovirus E2A DNA-binding protein was visualized in the same manner using dilute hybridoma culture supernatant fluid from the mouse monoclonal antibody clone B6-8, kindly provided by A. Levine of Princeton University (56). The exposed HyBlot ES Autoradiography film was scanned and the optical density of the specific signal quantified with the tools available in ImageJ.

### Chromatin immunoprecipitation (ChIP) assay

A ChIP assay was performed as previously described (57) on cells approximately 1 month post infection. Briefly, cells were cross-linked with 1% formaldehyde for 8 min at room temperature. Crosslinking was stopped by the addition of 0.125 M glycine for 5 min at room temperature. Cells were lysed using cell lysis buffer (5 mM PIPES pH 8, 85 mM KCl, 1% igepal) and protease inhibitors for 15 min on ice. The cell lysate was centrifuged at 2100 rpm for 5 min at 4°C. The supernatant was discarded and the pellet was resuspended in SDS lysis buffer (1% SDS, 10 mM EDTA, 50 mM Tris pH 8.0, dH2O) and protease inhibitors for 25 min on ice followed by flash freezing in liquid nitrogen. Lysed nuclei were sonicated using a Bioruptor water bath sonicator for 15 sec “On” and 30 sec “Off” 3 times to generate an average of 800 bp of sheared DNA, from which the 15 ul input sample was taken. The sonicated lysates were pre-cleared with salmon-sperm coated agarose beads (Upstate) and lysates were divided equally. One half of the lysate was immunoprecipitated with 5 µg of ChIP-grade antibody to RUNX1 (Abcam ab23980) overnight at 4°C. The other half of the lysate was immunoprecipitated with isotype control antibody. Immunoprecipitated proteins were incubation with 60 µl of salmon-sperm coated agarose beads for 2 h and then washed for 3 min at 4°C with the following buffers: low salt buffer (0.1% SDS, 1% Triton X-100, 2 mM EDTA, 20 mM Tris pH 8.0, 150 mM NaCl, dH2O), high salt buffer (0.1% SDS, 1% Triton X-100, 2 mM EDTA, 20 mM Tris pH 8.0, 500 mM NaCl, dH2O), LiCl buffer (0.25 M LiCl, 1% NP40, 1% DOC, 1 mM EDTA, 10 mM Tris pH 8.0, dH2O) and 1X TE buffer. DNA was then eluted with SDS elution buffer (1% SDS, 0.1 M NaHCO3, dH2O). After DNA elution, crosslinking was reversed overnight with 5 M NaCl at 65°C followed by treatment with proteinase K for 1 h at 45°C. Immunoprecipitated DNA was isolated using a phenol:chloroform:isopropanol mix (Invitrogen) as per the manufacturer’s instructions. QPCR reactions were run in triplicate on an ABI prism 7900 (Applied Biosystems, Foster City, CA). Isolated DNA was analyzed by real-time PCR using the following primers to E3, hexon, or GAPDH: E3 promoter region: (Sense sequence, 5’-CCCGCTCCCACCACTGT-3’, anti-sense sequence, 5’-TGCGCCCCTGAGTTAGTCA-3’, probe sequence, 5’-6 FAM-CCCAGAGAC-ZEN-GCCCAGGCCG-3IABkFQ-3’).

Hexon gene region (Sense sequence, 5’-GCCATTACCTTTGACTCTTCTGT-3’, anti-sense sequence,5’-CCTGTTGGTAGTCCTTGTATTTAGTATC-3’, probe sequence, 5’-6 FAM-AGAAACTTCCAGCCCATGAGCCG/36-TAMSp/-3’).

GAPDH gene region (Sense sequence, 5’-AAATGAATGGGCAGCCGTTA-3’, anti-sense sequence, 5’-TAGCCTCGCTCCACCTGACT-3’, probe sequence, 5’-FAM-CCTGCCGGTGACTAACCCTGCGCTCCT-QSY7-3’)

Values from real-time PCR reactions were analyzed using the ΔΔCt method, using the gene of interest input values to normalize samples (which were run alongside the ChIP sample DNA), followed by normalizing to the negative control isotype antibody sample, which was set to 1.

## Results

### The acute phase of adenovirus infection can proceed in B cells expressing *RUNX1*-containing leukemic fusion genes

Several RUNX1 proteins disrupt the organization of intranuclear sites of viral RNA synthesis (44). We therefore postulated that RUNX1-related leukemic fusion proteins may inhibit adenovirus infection in lymphocytes by interfering with viral replication. To evaluate the impact of the leukemic fusion genes on the course of an adenoviral infection, we created stable B cell lines expressing either the *RUNX1* fusion gene commonly associate with ALL (*ETV6/RUNX1*), or the *RUNX1* fusion gene commonly associated with acute myeloid leukemia (*RUNX1/MTG8*) (58, 59). While both fusion proteins can repress gene expression (39, 41, 60), the N-terminus of the *RUNX1/MTG8* fusion protein is derived from the RUNX1 protein and differs from the N-terminus of the *ETV6/RUNX1* fusion. Because the N-terminus of RUNX1 was shown to interact with adenoviral proteins and affect their localization (44), we reasoned that the fusion products may have different effects on the virus infection. For these experiments, cells were transfected with the indicated fusion genes or an empty vector and selected for several weeks to create stable cell lines.

The *ETV6/RUNX1* and *RUNX1/MTG8* fusion product detected by RT-PCR in stably transfected B cell lines was indistinguishable from that detected in transiently transfected A549 epithelial cells (**Fig 1**). The patient-derived UoC-B4 cells served as a positive control for the *ETV6/RUNX1* translocation (61). KASUMI-1 cells, derived from an AML, served as a positive control for the expression of the *RUNX1/MTG8* fusion gene (62). Continued presence of the fusion gene transcript was confirmed by RT-PCR throughout the course of the experiments. Western blotting confirmed expected presence of both ETV6/RUNX1 and RUNX1/MTG8 fusion proteins as well as the wild-type RUNX1 isoforms (**Fig 2**). We used a RUNX1/MTG8 construct which is similar in size to the truncated variant lacking the NHR3 and NHR4 regions of MTG8 as previously described (63, 64). This variant protein associates with histone deacetylase complexes, but does not disrupt cell cycle (65). This variant has been detected in clinical isolates as well as Kasumi-1 cell line (64). All variants of the RUNX1 and RUNX-fusion proteins identified in Fig 2A were detected. Although the levels of the RUNX1/MTG8 variant fusion protein are lower than wild-type RUNX1 expression, the level of RUNX1/MTG8 variant fusion protein appear similar between the Kasumi-1 cells and that directed by the expression vector in our established cell line (**Fig 2B**). In contrast, there is much greater expression of ETV6/RUNX1 in our transfected cell line as compared to the UoC-B4 control cell line (Fig 2C), and fusion gene expression is also much greater than expression of wild-type RUNX1 in our established cell line. The use of a ChIP-grade antibody demonstrates similar levels of RUNX1b and RUNX1c isoforms across the three stably transfected cell lines (**Fig 2C**). This antibody is, however, unable to detect the RUNX1/MTG8 fusion protein which lacks the ab23980 epitope (**Fig 2A**).

**Figure 1.**
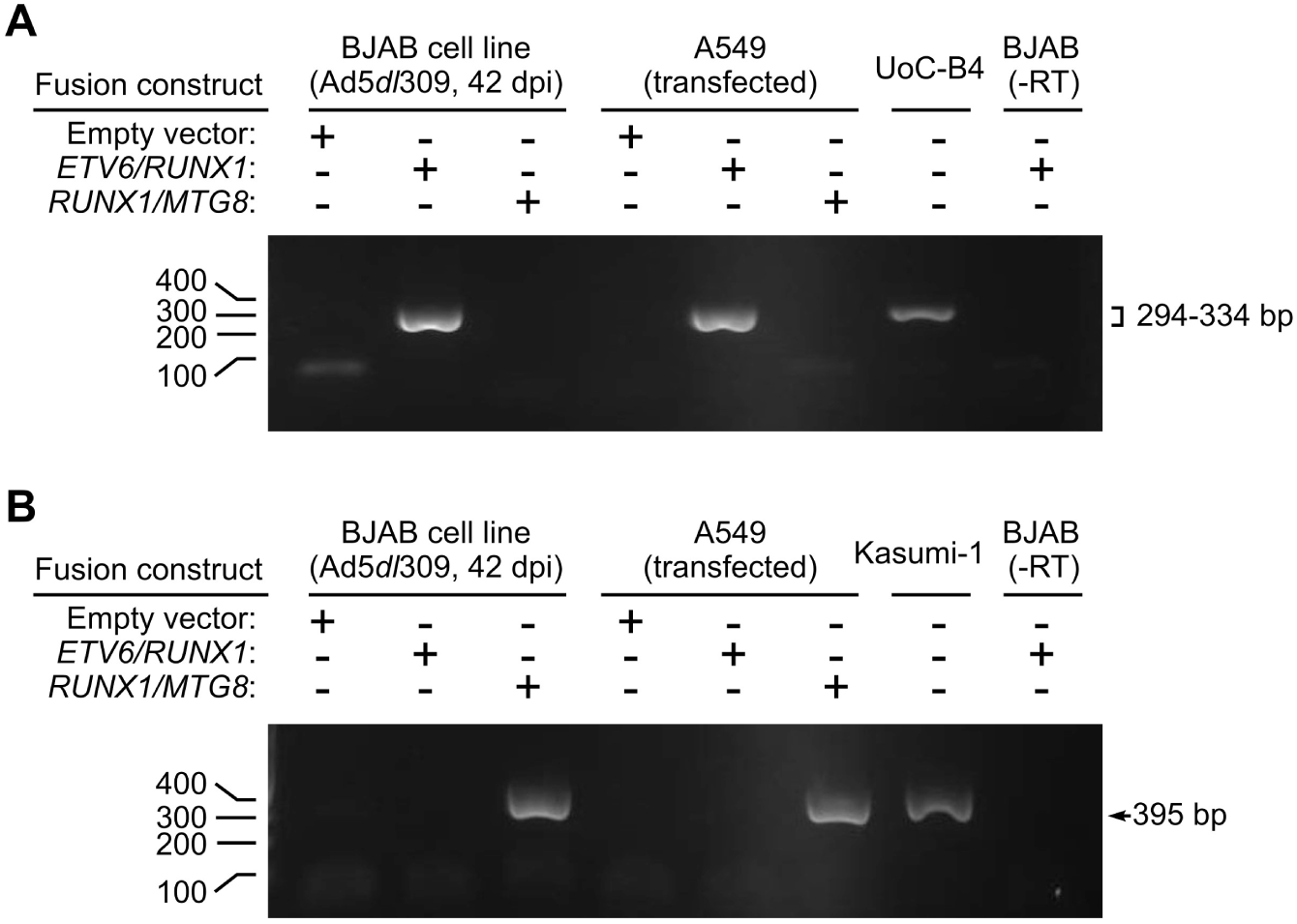
Stable expression of the ETV6/RUNX1 and RUNX1/MTG8 leukemia fusion genes persists in B lymphocytic cells following transfection and selection. BJAB cells were transfected with the pTARGET expression vector (Empty vector) or the same vector expressing either ETV6/RUNX1 or RUNX1/MTG8. Stable cell lines were established and selected as described in the Methods. mRNA was isolated from these cell lines 42 days after infection with Ad5dl309 and analyzed by PCR following reverse transcription. To serve as a control, RNA was isolated from A549 cells transiently expressing the same constructs as well as the leukemic cell lines UoC-B4 and Kasumi-1. (A) The TEL-H (ETV6) and AML1-G (RUNX1) primer pair are described in (5) and yields a 294-334 bp product for the ETV6/RUNX1 fusion transcript (upper panel). (B) The AML1-A (RUNX1) and ETO-B (MTG8) primers described in (54) yields a 395 bp PCR product for the RUNX1/MTG8 fusion transcript. RNA from infected BJAB cells transduced with ETV6/RUNX1 served as a negative control by leaving out reverse transcriptase (-RT).

**Figure 2.**
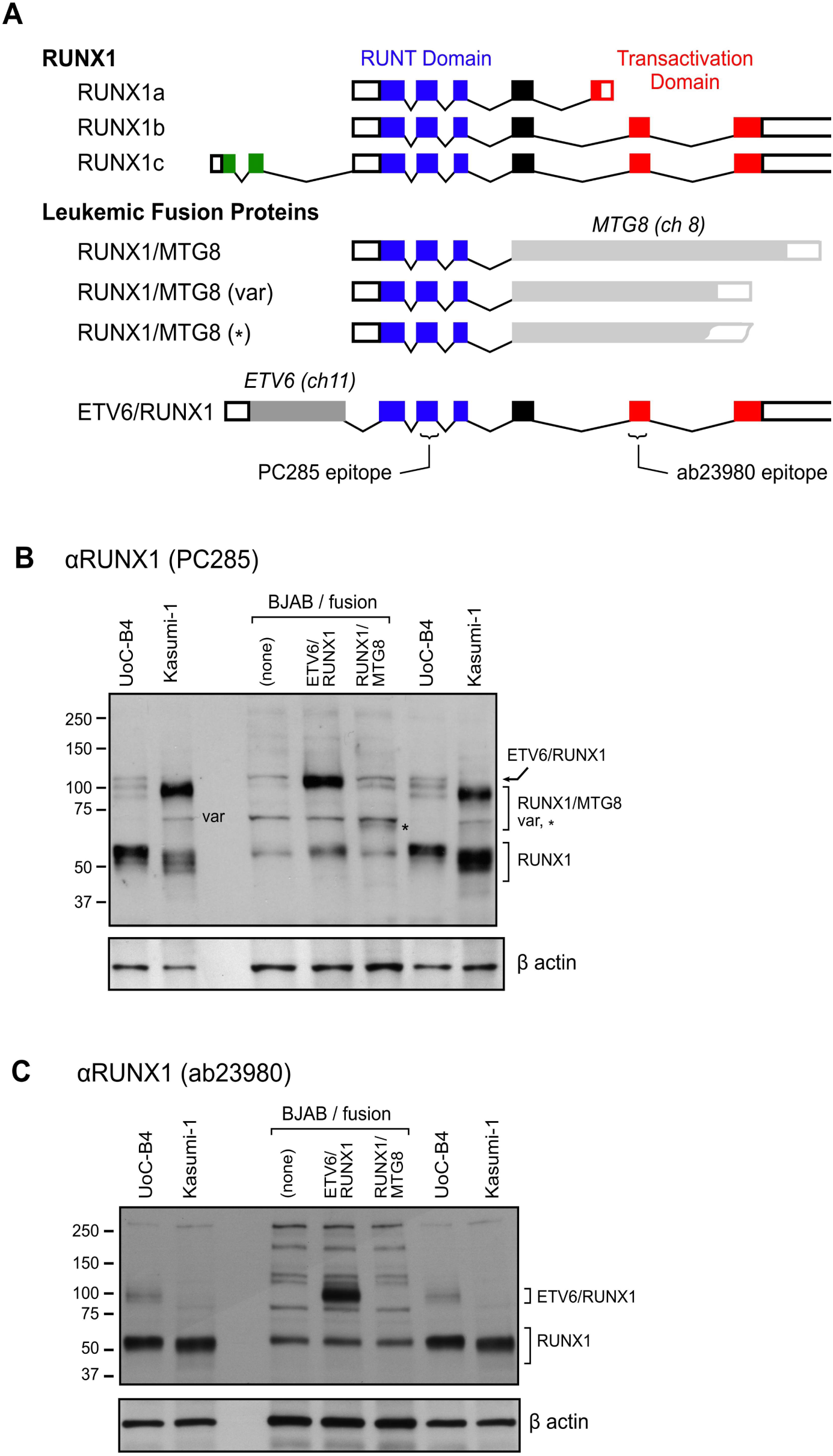
Schematic representation of the RUNX1 and RUNX1-related leukemic fusion proteins and RUNX1-antibody recognition sites. (A) The predominant isoforms of human RUNX1, the ETV6/RUNX1, and the RUNX1/MTG8 leukemic fusion genes are represented by boxes for exons where filled boxes are coding regions and open boxes are non-translated regions. Splices in the RUNX1 gene are indicated by segmented lines connecting the exons. The DNA-binding (RUNT) domain is shown in blue, the transactivation domain in RUNX1 in red and the unique exon incorporated into RUNX1c is green. The exon and intron structure of the fusion partners MTG8 and ETV6 are not shown. The predominant RUNX1/MTG8 fusion product (84kDa) is shown along with a naturally occurring variant (var) (62kDa). We used a RUNX1/MTG8 construct (*) which is similar in size to the truncated variant lacking the NHR3 and NHR4 regions of MTG8. The symbol “var” and “*” identify the likely RUNX1/MTG8 variant and variant construct in the corresponding immunoblot (B). Cellular lysates from UoC-B4, Kasumi-1 and stably transfected B lymphocytic cells expressing the RUNX1-related fusion genes were separated by electrophoresis through polyacrylamide with SDS, immobilized, and probed by immunoblotting with the RUNX1 antibody specific for the RUNT domain (clone PC285). (C) Immobilized cellular lysates were also probed with the ChIP-grade RUNX antibody clone ab23980 which detected the expected 100 kDa fusion protein in both UoC-B4 and transfected BJAB cells. This antibody detected the normal RUNX1b and RUNX1c isoforms (45-55 kDa) while RUNX1a isoform lacks the epitope recognized by the ab23980 antibody. β-actin was used as a control for protein loading.

Cells stably expressing the fusion genes were infected with serotype 5 adenovirus and viral protein expression was evaluated. A similar fraction of infected cells was found to synthesize a representative viral structural protein (hexon) whether transduced with the *ETV6/RUNX1, RUNX1/MTG8,* or an empty vector (**Fig 3A**). The levels and pattern of hexon expression measured by flow cytometry in these lymphocytic cells resembled that measured in acutely infected epithelial cells (66). The levels of hexon protein reached maximum levels (between 80-90% hexon-positive) on the same day post-infection (p.i.) in each of the cell lines evaluated (**Fig 3B**), however, between independent infections the levels of maximum hexon protein could vary between day 7 and 14 post infection as previously reported. Peak expression of hexon protein was similar between cells infected with either the Ad5dl309 (**Fig 3B**) or Ad5wt (**Fig 3C**) strain of the virus as expected since Ad5dl309 is phenotypically wild-type in cell culture since it only lacks proteins needed for immune evasion of host cells. The progress of the infection over the first 20 days resembled that previously described for lymphocytic cell lines (55) and we term this period as the “acute” phase of the infection. After 20 days of infection, hexon protein levels diminished and became undetectable by flow cytometry (55). We designate this phase of the infection as the “persistent” phase because the viral genome persists in several lymphocytic cell lines for over a year, during which time, viral late gene protein expression is often undetectable (55). Neither the ALL- nor AML-associated translocation appeared to alter the synthesis of hexon protein during the acute phase of the infection (Fig 3A-3C). The synthesis of hexon protein also decayed in a comparable manner between the samples; neither B cells containing the empty vector, nor those expressing the translocations, contained high amounts of intracellular hexon protein by day 21 (<15%, data not shown).

**Figure 3.**
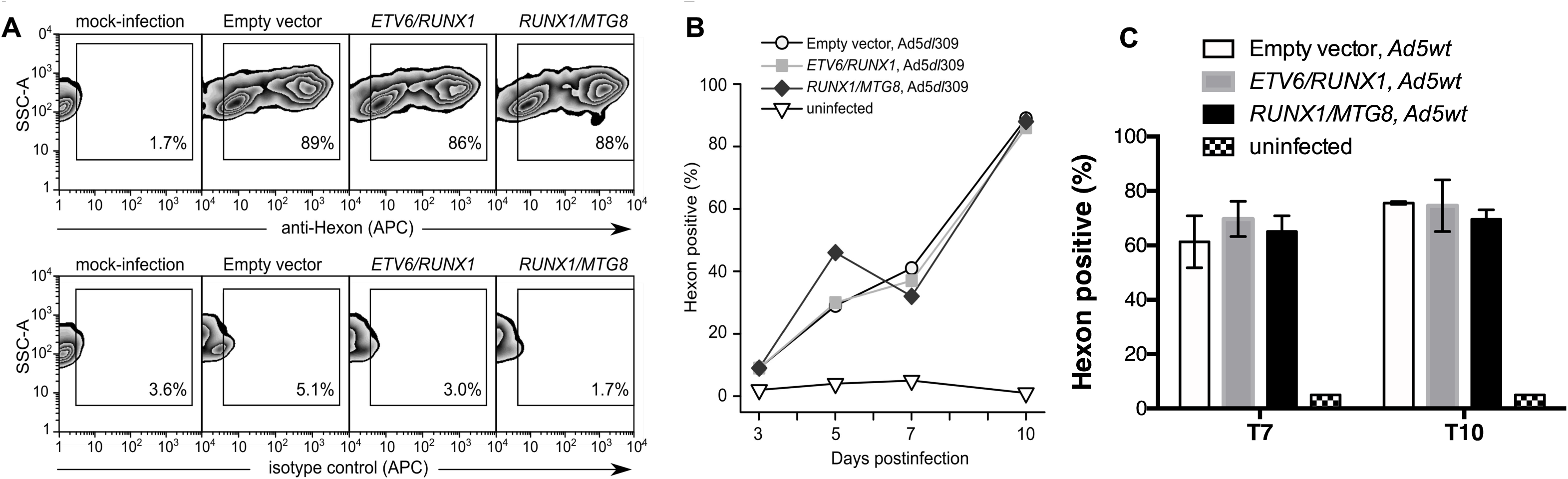
The “acute” phase of adenovirus infection in B lymphocytes is not impacted by enforced expression of the leukemia-associated ETV6/RUNX1 or RUNX1/MTG8 fusion genes. BJAB cell lines transduced with the empty (pTARGET) vector or the indicated fusion gene were infected with Ad5dl309 virus and stained at various days post-infection with a mouse monoclonal antibody to either hexon protein or a non-specific isotype control antibody followed by an APC-conjugated goat anti-mouse IgG antibody. (A) Representative flow cytometry plots showing infected cells stained for hexon (top panel) or with the isotype control (bottom panel) at 10 days post-infection. The fraction of hexon-positive cells is shown as a function of days post-infection in cells infected with either (B) Ad5dl309 or (C) Ad5wt.

### *RUNX1* containing fusion genes reduce adenoviral gene expression during persistent infected of B cells

Although hexon protein becomes undetectable during the persistent phase of the infection (55), hexon transcripts are detectable at much later times post-infection. Ten days p.i., the level of hexon mRNA is comparable to the level detected 3 days after infection (set to 1) and is similar between the cell lines evaluated (**Fig 4A**). However, 6 weeks later (day 54 p.i.), once a persistent infection is established, the amount of hexon mRNA detected is 4-logs lower in empty vector containing cells but became undetectable in cells containing either translocation. To determine more precisely when the loss begins to occur, we tracked cells over time in separate infections. B cells transfected with the empty vector and infected with Ad5 contain similar levels of hexon mRNA throughout the 7-week period shown in Fig. 4B. By contrast, levels of hexon mRNA declined over this period of time in cells expressing either the *ETV6/RUNX1* or *RUNX1/MTG8* fusion gene. One limitation of our experiments is the inherent variability between individual infections during the establishment of persistent infections in lymphocytes as compared to lytic infections in epithelial cells. Because of these differences, data from three independent infections are shown to demonstrate that a much greater loss of AdV hexon mRNA consistently occurs in the presence of either *ETV6/RUNX1* or *RUNX1/MTG8* fusion genes. While the length of time required for loss of hexon mRNA varies between three replicate infections, expression was robustly reduced in *RUNX1/MTG8* or *ETV6/RUNX1*-expressing B cells between 7 to 10 weeks post infection (**Fig 4A-4C**). Hexon expression remained detectable in the translocation-negative cells over this time period, though there was some variability in the amount of late AdV gene expression retained from infection to infection.

**Figure 4.**
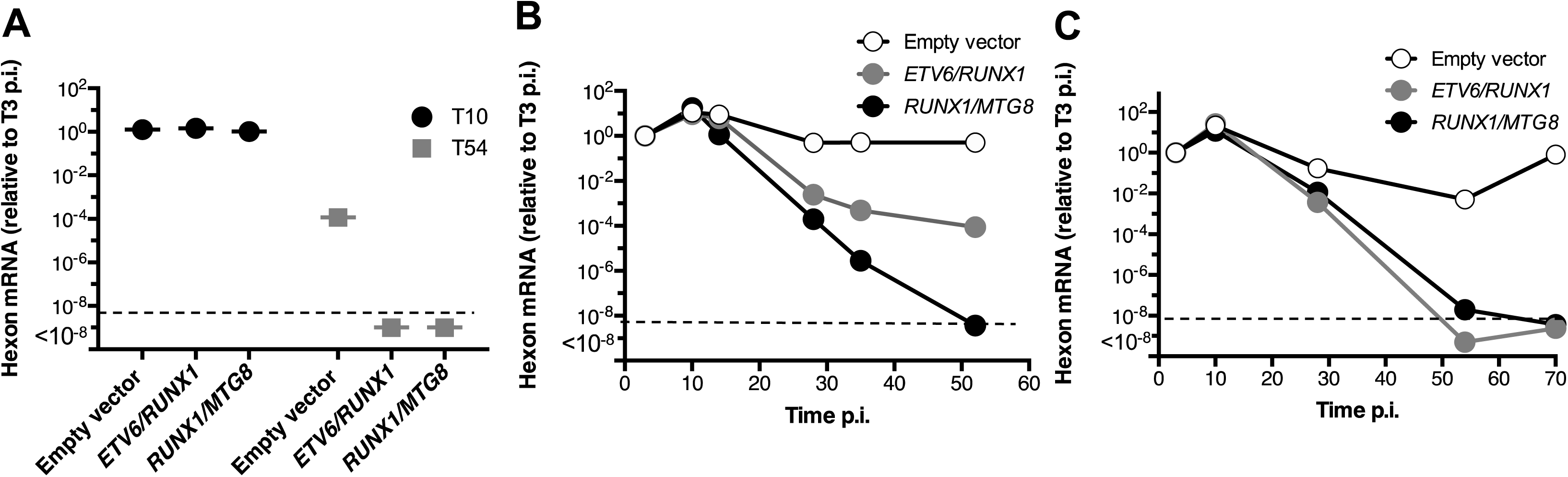
Expression of the late viral gene *hexon* declines in persistently infected B lymphocytes that contain the leukemia-associated ETV6/RUNX1 or RUNX1/MTG8 fusion genes. (A-C) BJAB cells stably expressing either the empty vector, ETV6/RUNX1 or RUNX1/MTG8 were infected with Ad5dl309. At the indicated times, the level of hexon mRNA was measured by RT-qPCR and normalized to EIF1 expression as described in *Material and Methods*. Samples are shown relative to the level of expression 3 days post-infection (which was set to 1). The dotted line represents the limit of detection for the qPCR (Ct=undetermined). T10 (10 days p.i.) and T54 (54 days p.i.). *Three independent infections are shown*.

To determine the extent of viral gene expression loss, the levels of representative early transcripts from three other transcription units were measured. Twenty-eight days into the infection, hexon, E2A and E3-gp19K mRNA levels were significantly reduced in cells expressing either *ETV6/RUNX1* or *RUNX1/MTG8*RUNX1 compared to expression levels in empty vector containing cell (**Fig 5A-D)**. *E3-gp19K* appeared to be reduced the most by the translocations and exhibited approximately 15% the expression detected in empty vector cells (**Fig 5D**). By comparison, the translocations caused expression of *hexon* to be between 20-25% of that seen in control cells (**Fig 5A**). In contrast, expression of E1A was much more variable and was not significantly reduced. Among the viral genes expressed during persistence, and in the absence of RUNX1 translocations, *hexon* expression is the highest. We compared the expression levels of *E1A-13S*, *E2A*, and *E3-GP19K* relative to *hexon* levels (solid line at 1) detected 28 days p.i. and then measured the magnitude of loss 2-weeks later (42 days p.i.) (**Fig 5E-G**). Control cells containing the empty vector did not exhibit a significant loss in expression of any of the early genes measured. In contrast, early gene expression in translocation-positive cells was significantly reduced at 42 days p.i., as compared to 28 days into the infection. Again, the levels of AdV early expression were different between the three independent infections 28 days post-infection. However, there was a reduction in early gene expression observable across independent infections with complete loss of all three genes observed by 6-weeks post infection in at least one infection of the three. These results suggest that the presence of the RUNX1-related fusion genes reduced expression of both early and late adenovirus genes in a manner that is independent of the sequences at the N-terminus or C-terminus of the RUNX1 protein. Moreover, this effect is likely to be independent of the RUNX1 transactivation domain, which is absent in the RUNX1/MTG8 fusion protein. Overall, in contrast to the similar pattern of viral protein expression observed among the cell lines during the acute phase of the infection, the *RUNX1*-related fusion genes altered the course of the infection during the persistent phase by reducing viral mRNA expression. Notably, the decrease in viral gene expression was not due to changes in the viability or growth of the infected cells which was unaffected by the presence of the *RUNX1*-related fusion gene (**Fig S1**).

**Figure 5.**
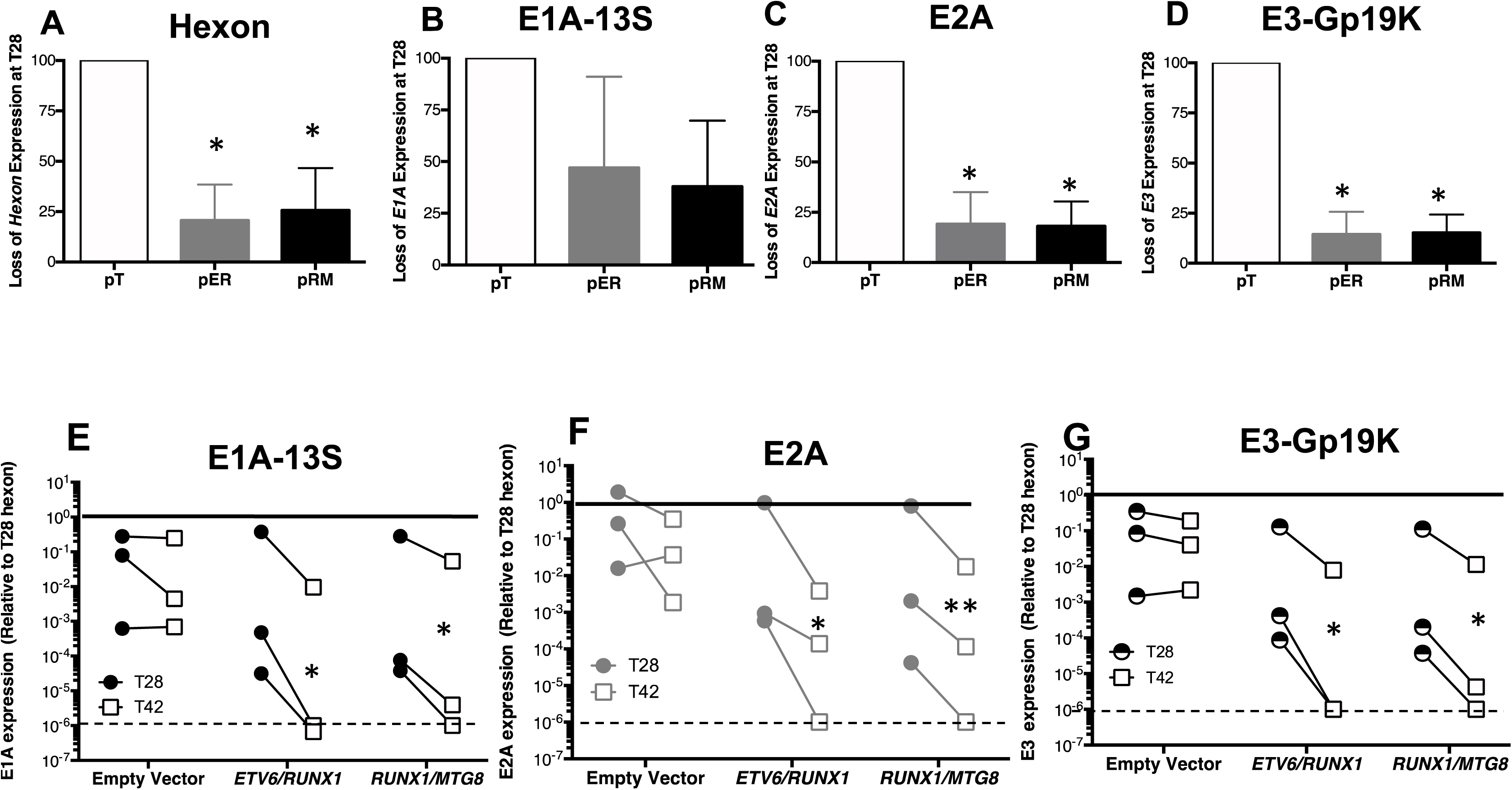
Expression of early viral genes is reduced in persistently infected B lymphocytes that contain the leukemia-associated ETV6/RUNX1 or RUNX1/MTG8 fusion genes. (A-D) BJAB cell lines stably transfected with constructs to express the indicated leukemia-associated genes were infected with Ad5dl309 and the levels of the indicated adenovirus genes were measured 28 days p.i. The level of expression of each gene in empty vector containing cells (pT) is set to 100% for comparison to the level in cells containing either ETV6/RUNX1 (pER) or RUNX1/MTG8 (pRM) fusion genes. *Data shown are the SEM of three independent experiments. * indicates P<0.05 of an unpaired student T-test.* (E-G) Expression of *E1A-13S*, *E2A* and *E3gp19K* were measured on days 28 (circles) and 42 (squares) post-infection. Samples were normalized to the housekeeping gene E1F1 and expression of each gene is shown relative to *hexon* expression for each sample at T28 (which is set to 1) for comparison between genes. *Data shown is from 3 independent experiments and significance of ratio paired T-test *P<0.05, **P<0.01*.

### Adenovirus genome is not retained in persistently infected B cells that express leukemic fusion genes

The loss of viral transcripts could reflect repression of transcription or loss of the viral genome. At the start of infection, we measured AdV DNA levels in cells containing the leukemic fusion genes compared to empty vector containing cells. Interestingly, though we detected almost no difference in AdV hexon protein levels or mRNA 10 days p.i., we detected a consistent increase in AdV genome copy number during acute infection in cells containing the *ETV6/RUNX1* or *RUNX1/MTG8* fusion gene (**Fig 6A**). Again, there was a considerable amount of heterogeneity in the copy number present during each infection but the fold change significantly increased 6-fold in ETV6/RUNX1-positive cells and 5-fold in RUNX1/MTG8-positive cells on average in each infection (**Fig 6B**). The amount of viral DNA in persistently infected BJAB cells stably transfected with the *RUNX1*-related fusion genes or empty vector was measured over the course of several weeks. Because we detected variable amounts of viral DNA at the peak of infection, we normalized AdV DNA levels measured over time to the level present at the start of each experiment. Results in **Fig. 6C** show the changes in viral DNA relative to the amount present at the initial peak of infection (10 to 14 d post-infection) for three independent experiments. Levels of viral DNA dropped in all three infections of cells transduced with the empty vector but remained relatively stable around 40 days after infection. By contrast, loss of the viral genome was greatly accelerated in cells expressing either the *ETV6/RUNX1* or *RUNX1/MTG8* fusion gene. B cells containing either fusion gene exhibit a 6- to 8-log reduction in viral genome levels from peak levels. As the infection progressed, only a small fraction of the amount of viral DNA was retained in RUNX1 fusion containing cells as compared to translocation negative samples at the same time post infection. For example, one infection contained 2.2 x a10^4^ AdV genomes in *ETV6/RUNX1*-containing cells, 52 days post infection, as compared to 1.3 x 10^8^ in empty vector containing cells at the same time (**Fig 6D**). *RUNX1/MTG8*-containing cells contained 2.7 x a10^3^ AdV genomes. In another infection, AdV DNA levels were 7.9 x a10^6^ , 430, and 1337 in empty vector, *ETV6/RUNX1*-, and *RUNX1/MTG8*-containing cells, respectively 56 days into the infection. In comparison, the infection exhibiting the large reduction in viral DNA in translocation-negative cells over time still retained approximately 7,200 copies of viral genome per 100 cells at 62 days post-infection.

**Figure 6.**
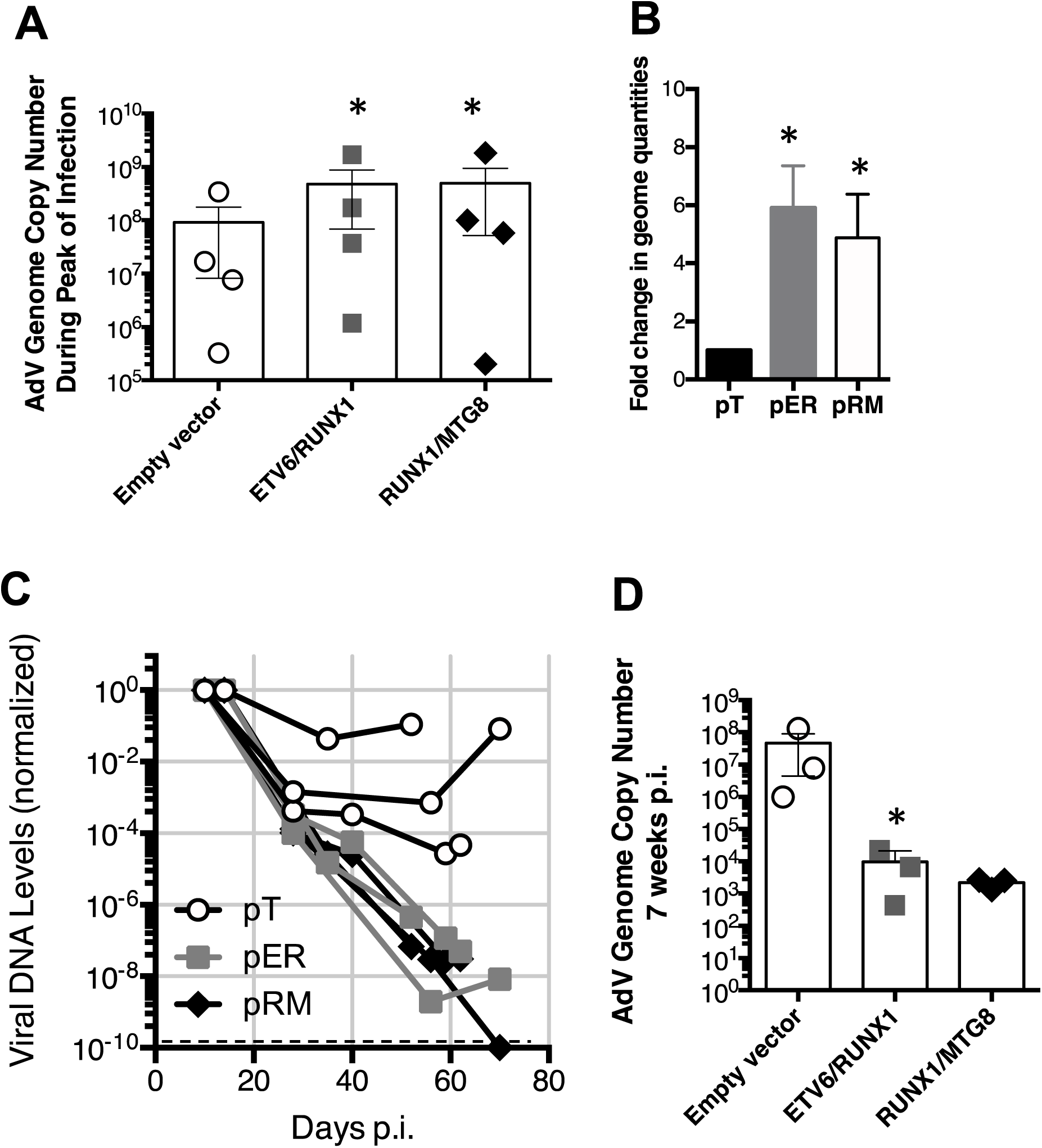
Expression of ETV6/RUNX1 or RUNX1/MTG8 does not reduce initial levels of viral genome in infected B lymphocytes but promotes the loss of adenoviral DNA during persistence. (A) AdV DNA levels were quantified by standard curve qPCR as described in *Material and Methods.* AdV genome copy numbers detected during acute infection (days 13-17 p.i.) with Ad5wt and normalized to GAPDH DNA are shown. Bars represent SEM of four independent experiments. * indicates P<0.05 of Pearson correlation coefficient between pairs infected at the same time (empty vector vs ETV-RUNX1 or empty vector vs RUNX/MTG8). (B) The fold-change in genome copy levels detected in cells containing ETV6/RUNX1 (pER) or RUNX1/MTG8 (pRM) over those detected in the empty vector (pT). Statistical significance determined by an un-paired T-test of the SEM of fold change from four experiments. *\*=P<0.05*. (C) BJAB cells stably expressing either an empty vector (pT; open circle) or the indicated leukemia-associated fusion gene ETV6/RUNX1 (pER; grey box) or RUNX1/MTG8 (pRM; black diamond) were infected with Ad5dl309. On the indicated days post-infection, viral DNA levels were measured by qPCR and normalized to the level of the cellular gene GAPDH. For each independent infection, viral DNA levels are shown relative to the level detected during acute infection (10-14 days p.i.); set to 1. The dotted line represents the limit of detection for the qPCR (Ct=undetermined). (D). AdV DNA levels were quantified 6-7 weeks p.i. (T52-59) and normalized to cellular levels of GAPDH DNA. SEM of 3 independent infections. * indicates P=<0.05 of a ratio paired T-test.

The Ad5dl309 virus used in some of these, and previous (55), experiments contains a deletion of the 14.7K, 14.5K, and 10.4K proteins within the E3 region of the viral genome. We next wanted to determine if there was an interaction between RUNX1 fusion genes and the viral genome that could be contributing to the loss of viral genome. To ensure that all of the viral DNA was present for these analyses we utilized Ad5wt in a separate set of experiments. We first confirmed the loss of viral DNA using the wild-type viral strain. Interestingly, cells transduced with the empty vector and infected with Ad5wt, with a replete E3 region, retained a more constant level of the viral genome over the entire five-week observation period (**Fig 7A**) when compared to infections with Ad5dl309 (**Fig 6B**). However, Ad5wt-infected cells expressing *ETV6/RUNX1* still suffered as much as a 4-log loss of viral DNA over this observation period. The rate of genome loss for Ad5dl309 (**Fig 5C**), however, was substantially and significantly greater (p < 0.04 by log-linear regression) than the loss of the Ad5wt genome in cells containing the fusion gene. Thus, these data reveal an unexpected difference between the true wild-type Ad5 and phenotypically wild-type Ad5dl309 virus.

**Figure 7.**
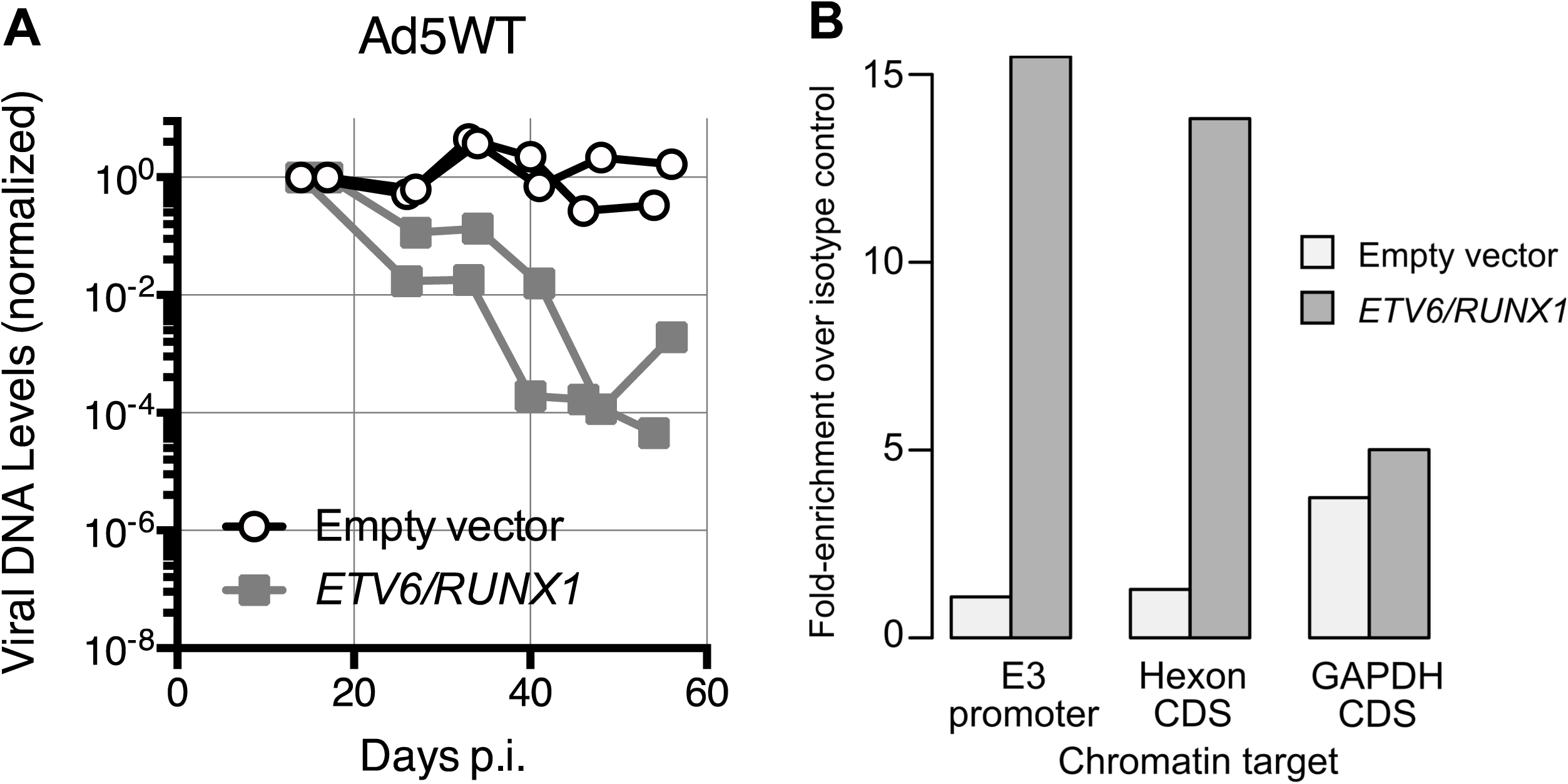
The leukemic associated fusion protein ETV6/RUNX1 binds the adenoviral genome. (A) ) BJAB cells stably expressing either an empty vector (open circle) or the leukemia-associated fusion gene ETV6/RUNX1 (grey box) were infected with Ad5wt. On the indicated days post-infection, viral DNA levels were measured by qPCR and normalized to the level of the cellular gene GAPDH. For each independent infection, viral DNA levels are shown relative to the level detected during acute infection (10-14 days p.i.) which was set to 1. Results are shown for two independent experiments. (B) BJAB cells stably expressing either an empty vector or ETV6/RUNX1 were infected with Ad5wt. On days 24 and 31 post-infection, cells were fixed and chromatin immunoprecipitation was performed with either a non-specific isotype-matched antibody as a control or an antibody specific for the RUNX1 protein (ChIP-grade RUNX antibody clone ab23980). DNA that immunoprecipitated with the antibodies was measured by qPCR with primers specific for the promoter region of the adenovirus E3 transcription unit (contains two RUNX1 binding sites) and the adenovirus *hexon* gene coding region (contains one RUNX1 binding site), or the coding region of the cellular GAPDH gene (no RUNX1 binding sites). The total amount of DNA (input) used for the immunoprecipitation was determined using 3% of the total lysate and used to normalize the amount of specific DNA recovered. Samples were further normalized to the amount of DNA immunoprecipitated with the isotype control antibody. *The average of two independent experiments are shown*.

### The ETV6/RUNX1 fusion protein binds adenovirus DNA

RUNX1 fusion proteins repress gene expression by directly binding chromatin and recruiting transcriptional repressors (39). To determine if the ALL associated RUNX1-fusion protein bound the viral genome, the E3 promoter region and the *hexon* coding region, which both contain RUNX1 binding sequences, were queried by a chromatin immunoprecipitation (ChIP) assay. *RUNX1/MTG8* expressing cells were not evaluated because the RUNX1/MTG8 fusion protein lacks the RUNX1 epitope recognized by the ChIP-grade antibody used for these experiments (**Fig 2A and 2C**). B cell lines stably expressing either the *ETV6/RUNX1* fusion gene or the empty vector were infected with adenovirus and evaluated for RUNX1 binding to viral DNA just before the establishment of viral persistence, and prior to significant loss of viral DNA by the fusion protein (24 and 31 days post-infection) (**Fig 7B**). More viral DNA was immunoprecipitated by the RUNX1 antibody from cells expressing the *ETV6/RUNX1* fusion gene than from cells transduced with the empty vector. Immunoprecipitation of DNA required a canonical RUNX1-binding site because a region of the cellular *GAPDH* gene devoid of RUNX1-binding sites within 800 nucleotides of the target sequence was not recovered in excess from *ETV6/RUNX1*-expressing cells. Because both empty vector-transduced and *ETV6/RUNX1*-transduced cell lines express similar amounts of endogenous RUNX1 protein (**Fig 2C**) the additional RUNX1-related protein recovered by chromatin immunoprecipitation is most likely the ETV6/RUNX1 fusion protein.

### HDAC inhibitors increase viral gene expression in *ETV6/RUNX1*-expressing cells

The chromatin-bound ETV6/RUNX1 proteins can recruit HDACs to repress transcription (39), so we were curious to see if HDAC inhibition with Trichostatin A (TSA) could increase viral mRNA in persistently infected B cells expressing *ETV6/RUNX1*. We first, evaluated what impact inhibition of HDACs would have on AdV gene expression in the absence of the repressive fusion gene and were surprised to see that treatment with TSA increased expression of all the viral genes evaluated to some degree in persistently infected cells (**Table 1; Empty vector**). *E1A* mRNA and *E3* mRNA increased 4- and 7-fold, respectively, in infected cells treated with TSA. *Hexon* mRNA increased robustly by 28-fold after treatment. De-repression of *E3,* however, was greater in TSA-treated cells expressing *ETV6/RUNX1* (16-fold) than in cells containing the empty vector. The magnitude of change in *E2A* mRNA (8.2-fold empty vector vs 15.8-fold ETV6/RUNX1) upon HDAC inhibition was similar to that detected in E3. Interestingly, both E2A and E3-gp19K expression appeared to be slightly more repressed in translocation containing cells that either E1A or hexon expression 28 days into the infection (Fig 5A-D). Overall, *E1A* levels exhibited the lowest response to inhibition of HDACs with a 4- and 5-fold increase following TSA treatment in empty vector-transduced and *ETV6/RUNX1*-transduced cells, respectively. These results may indicate that some viral genes are more strongly repressed by HDACs than others (*E3* and *hexon*) and even more so in the presence of the ETV6/RUNX1 fusion protein (*E3* and *E2A*).

**Table 1.** HDAC inhibition increases viral gene expression in persistently infected cells.

| Leukemic fusion<br>gene | Viral mRNA |  |  | Viral<br>DNA |
| --- | --- | --- | --- | --- |
|  | E1A | E3gp19K | Hexon |  |
| Empty vector | 4 ± 2 | 7 ± 1 | 28 ± 13 | 2 ± 1 |
| *ETV6/RUNX1* | 5 ± 2 | 16 ± 1 | 29 ± 7 | 9 ± 3.5 |

B cells stably expressing either the empty vector or the leukemic fusion gene *ETV6/RUNX1* were mock-infected or infected. Persistently infected cells were treated with the 175 nM TSA or vehicle control 30-40 days p.i. After 48 h, viral genomic DNA was quantified by qPCR and the indicated viral mRNAs were quantified by qPCR after reverse transcription. Values shown represent the fold increase after TSA treatment over untreated cells. *Data represents SEM of 3 independent experiments*.

### Viral DNA-binding protein amounts decline rapidly in *ETV6/RUNX1* expressing cells and precedes the decline in the viral DNA genome

Our data support the idea that ETV6/RUNX1 most likely represses E3/E2A expression by binding the viral genome (Fig 7B) and recruiting cellular histone deacetylases. Since the RUNX1-binding site at the E3 promoter is near the E2A promoter, any ETV6/RUNX1 bound to this site could also repress transcription from the E2A promoter located on the reverse strand. E2 encodes for the viral DNA-binding protein (DBP) and abundant levels of this protein are required for viral DNA replication (67). Consequently, a decline in E2A-DBP level would be expected to cause a corresponding decline in viral DNA levels. Interestingly, HDAC inhibition also increased the level of viral DNA more robustly in cells expressing the *ETV6/RUNX1* fusion (9-fold) compared to cells transduced with the empty vector (2-fold) (**Table 1**). We therefore compared the amount of E2A-DBP measured by immunoblotting with levels of viral DNA measure by qPCR at various times after infection with the wild-type Ad5wt virus, which retains higher levels of viral DNA over time (**Fig 7A**). In independent experiments, the detection of E2A-DBP remained stable between day 17 to 34 p.i. in cells transduced with the empty vector (**Fig 8A and 8B**). By sharp contrast, levels of E2A-DBP declined at an exponential rate in cells that expressed the *ETV6/RUNX1* fusion gene compared to translocation negative cells (**Fig 8C-D**). Moreover, for both cell lines, the decline in E2A-DBP levels preceded the decline in viral DNA levels (black circles). Although a less extensive time course was evaluated for Ad5dl309-infected cells, E2A-DBP again became undetectable in ETV6/RUNX1-expressing cells before becoming undetectable in cells transduced with the empty vector (day 21 in **Fig 8E**). It seems reasonable that the diminished level of E2A-DBP led to a decline in viral genome levels, and that the decline in viral genome levels is accelerated by the presence of the *ETV6/RUNX1* translocation.

**Figure 8.**
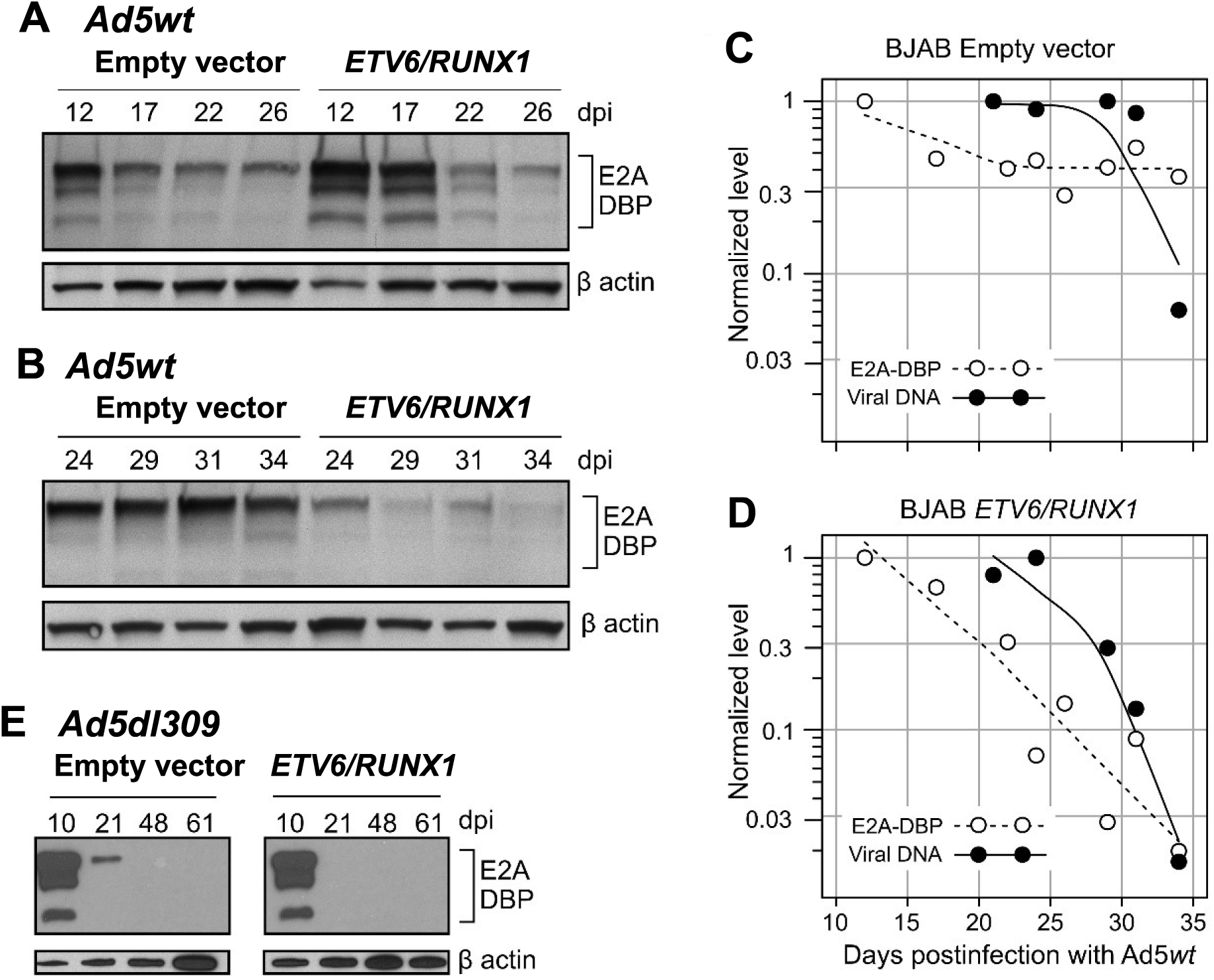
Levels of the viral protein required for viral DNA replication decline rapidly and in advance of the decline in viral DNA levels. BJAB cells stably expressing the empty vector or ETV6/RUNX1 were infected and evaluated by immunoblot for the viral E2A DNA-binding protein (DBP) and by qPCR for the viral DNA genome. (A, B) Cellular lysates from cells infected with Ad5wt were prepared on the days indicated and probed with antibodies against viral E2A-DBP and cellular β-actin protein. Levels of the proteins were detected by chemiluminescent imaging. (C-D) E2A-DBP and β-actin were quantified from the optical density of the chemiluminescent images shown in panels A and B with ImageJ. The level of E2A-DBP staining (open symbols) was normalized to that of β-actin and then compared to the level detected on the first day of evaluation (set to 1). Levels of viral DNA (closed symbols) from the corresponding infection were normalized to the level of the cellular GAPDH gene and compared to the maximum amount detected (set to 1). Local polynomial smoothing curves were generated from the entire span of data with the loss function in R. (E) Cellular lysates from cells infected with Ad5dl309 were prepared and analyzed as in panels A and B.

### Virus-induced repression of some cellular genes is retained after loss of the virus from *RUNX1* fusion gene expressing cells

We previously reported that expression of several cellular genes, including *CXADR, BNIP3, SPARCL1, SLFN11, BBS9* and *BNIP3,* is significantly downregulated by epigenetic means during the persistent phase of infection of B cells (17). Because of the durable nature of epigenetic changes, we wondered if these changes could remain in cells that no longer harbored the virus following expulsion by *ETV6/RUNX1*. Cells expressing the *RUNX1* fusion gene, or cells stably transfected with the empty vector, were infected with adenovirus and evaluated for repression of cellular genes at late times post-infection when most of the virus was lost from the population. Among the cellular genes repressed by the virus, *CXADR*, *SPARCL*, *SLFN11* and *BNIP3* were significantly de-repressed following treatment with epigenetic enzyme inhibitors suggesting that repression of these four genes may be sustainable. After two months of infection, the number of viral genomes was determined and the relative level of mRNA for *CXADR, BNIP3*, *SPARCL1* and *SLFN11* was measured in infected and mock-infected cells. As expected, viral DNA was retained in the persistently infected B cells without the translocation, which contained an average of 1.4 x 10^4^ genomes per 100 cells (**Table 2**). As previously shown, the viral genome was lost from infected B cells that harbored the leukemic translocations and cells expressing the *ETV6/RUNX1* translocation retained less than 2 genomes per 100 cells.

**Table 2.**
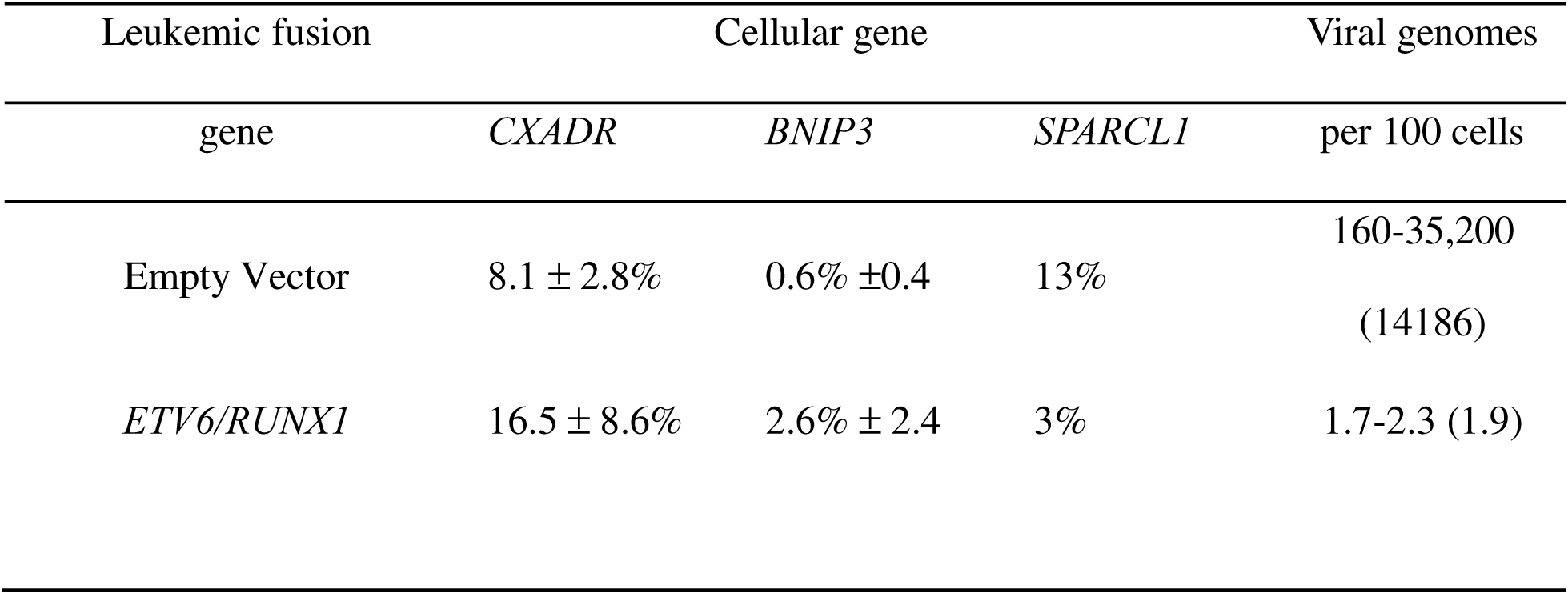
Adenovirus-mediated downregulation of cellular genes is retained after expulsion of the virus.

In agreement with previously reported results, infection reduced *CXADR, BNIP3,* and *SPARCL1* expression to between 0.6% and 13% of the level detected in non-infected cells transduced with the empty vector. Expression of these cellular genes remained repressed between 3% to 17% the level of that detected in non-infected cells, even in cells expressing the *ETV6/RUNX1* fusion gene. This reduction is striking in *ETV6/RUNX1*-expressing cells because at least 98% of these B cells no longer contained the viral genome. Because these cells contained only 1.7-2.3 genomes per 100 cells, it seems likely that the expression of cellular genes remained low in a subset of cells that had lost the viral genome. Expression of *CXADR, BNIP3,* or *SPARCL1* was not downregulated by the *ETV6/RUNX1* fusion in the absence of a viral infection (data not shown). Curiously, repression of *SPARCL1* by the virus was observed inconsistently in three separate infections. In the experiment where repression by the virus was detected (**Table 2**), however, the repression was retained after loss of the virus from the cells. In addition, we did not observe repression of *SLFN11* expression in our infected cells (either with or without the translocation) thus we could not evaluate its stable repression once the virus had been kicked out of cells. These findings highlight one important technical difference between our previous experiments and the current study which is that the cells used here are all under selection to maintain expression of the vectors. It seems likely that infected cells cell growing under antibiotic selection may not repress expression of cellular genes as they would under normal cell growth conditions. Nonetheless, repression of two of four cellular genes known to be repressed by the virus is consistently retained after the virus is kicked out of the cells. These results indicate that adenovirus may be capable of leaving an epigenetic echo of its infection in the form of durable epigenetic repression of cellular genes after the virus is expelled from the infected cell.

Levels of *CXADR, BNIP3,* and *SPARCL1* mRNA were measured during the persistent phase of infection with Ad5*dl*309 (56-77 days p.i.) and normalized to the housekeeping gene *EIF1*. The relative level of gene expression in infected B cells is expressed as a percentage of the level detected in uninfected cells (which is set to 100%). At the same time, the number of viral genomes was determined by qPCR (average of 3 experiments in parenthesis). *CXADR value mean from 3 independent experiments and BNIP3 from 2 independent experiments*.

## Discussion

Viruses are responsible for over twenty types of cancer and as much as 15% of the world-wide cancer burden (recently reviewed in (68, 69)). An infectious etiology for ALL has long been postulated (30, 70–72) and although a recent study suggests that congenital cytomegalovirus infection may be a risk factor for ALL (73) no microorganism has definitively been linked to this disease. One hypothesis that accounts for the absence of any link is that ALL associated mutations, induced by a common virus in the leukemic progenitor cell, preclude retention of the virus. The DNA tumor viruses JC virus (70) and adenovirus (28, 29) have been suggested to be a candidate for this common virus. In this study, we sought to evaluate the potential for mutations associated with the initiation of leukemia to perturb adenovirus infection by examining the impact of RUNX1 leukemic fusion genes on adenovirus persistence in B lymphocytic cells.

The program of adenovirus gene expression differs in lymphocytic cells during the acute phase, which is characterized by robust viral gene expression, and the persistent phase in which viral gene expression is gradually dampened. The common RUNX1 leukemic fusion genes *ETV6/RUNX1* and *RUNX1/MTG8* had no impact on viral gene expression during the acute phase but accelerated the decrease in viral gene expression and viral DNA replication during the persistent phase of infection. This decrease in viral gene expression was measured as a decrease in the levels of representative early and late viral mRNAs. The decrease in viral DNA levels was preceded by an accelerated loss of E2A DNA-binding protein in *ETV6/RUNX1* expressing cells. Since the E2A DNA-binding protein is essential for viral DNA replication (74, 75), reduced levels of this protein would logically lead to diminish viral DNA replication. We infer that viral DNA replication was affected because without viral DNA replication, viral genomes would have been diluted below the limit of detection (5-orders of magnitude) after less than four weeks (Fig. S1). We also cannot exclude the possibility that the stability of viral DNA was further decreased by the RUNX1 leukemic fusion genes.

An unexpected finding was that viral genome levels declined at a greater rate in cells infected with the E3-deleted virus Ad5dl309 than the wild-type virus (p < 0.04 by log-linear regression). E3 products promote immune evasion, confer resistance to death-promoting cytokines, and promote cell death at late times of a productive infection (76–78). Although these activities would impact the persistence of infected cells, it seems unlikely that the E3 14.7K, 14.5K and 10.4K proteins, which are missing in Ad5dl309-infected cells, directly control viral DNA replication. Nonetheless, expression of the E3 and E2A genes may be linked. The E3 14.5K and 10.4K protein have been reported to affect E1A expression (79), which would in turn affect E2A transcription. The E2A and E3 promoters direct transcription in opposite directions from a common position on the viral chromosome and similar levels of RNA pol II pre-initiation complex are found between these promoter regions (80). Because the E3 promoter can respond to activation signals independently of E1A (81, 82), co-stimulation of the E3 and E2 promoter may promote adenovirus persistence in lymphocytes where E2 products direct viral DNA replication while E3 products allow for immune evasion. These results may hint at additional roles for the E3 region in persistence of viral DNA even in the absence of immune pressure.

Viral gene expression appears to be repressed by epigenetic changes in persistently infected cells even in the absence of the ETV6/RUNX1 fusion protein. The adenovirus chromosome acquires histones shortly after entering the nucleus in absence of DNA replication (83, 84). This would render it susceptible to regulation by histone modifications, and acetylated histones have recently been identified at various viral promoters (80). Accordingly, viral mRNA levels increased after exposure to an HDAC inhibitor. The magnitude of E1A regulation by histone acetylation, or TSA-sensitive processes, is less substantial than that detected in the other viral genes evaluated. Additionally, since the inhibitor increased levels of E3gp19K more in ETV6/RUNX1-expressing than in vector-transduced cells (Table 1), the ETV6/RUNX1 fusion protein appears to repress specific adenoviral genes differentially. Both ETV6 and the ETV6/RUNX1 fusion proteins are site-specific DNA-binding proteins that recruit chromatin remodeling proteins to repress target gene transcription (39, 85, 86). The ETV6/RUNX1 protein was enriched at a site in the E3 promoter region as well as at a site distant from any promoter in the hexon coding region. Curiously, when considering empty vector cells, ChIP analysis showed no enrichment of endogenous RUNX1 on the viral chromosome (with canonical RUNX1 binding sites) over that seen at a cellular gene (with no RUNX1 binding sites). Since there was plenty of endogenous RUNX1 protein in translocation negative cells (Fig. 2C), endogenous RUNX1 may not bind the viral genome with the same avidity as the ETV6/RUNX1 fusion protein. Alternatively, interactions between other adenovirus proteins and the RUNX1 proteins (44) may limit access of the endogenous RUNX1 protein to the viral genome. These results suggest that RUNX1 fusion proteins can evict adenovirus in this B-cell model by histone deacetylation likely causing a subsequent loss of viral gene expression, that is followed by a decline in viral DNA replication.

We previously identified a small number of cellular genes that are downregulated in B-cells persistently infected with adenovirus (17). Here, we found that expression of three of these genes, *BNIP3, SPARCL1* and *CXADR*, remained repressed after the virus was evicted by ETV6/RUNX1 (Table 2). These results indicate that epigenetic remodeling of the cellular genome by adenovirus persists in the absence of the virus. This epigenetic echo could be part of a mechanism that contributes to hit-and-run transformation. We previously reported that *SPARCL1* and *CXADR* are downregulated in cell lines that harbor translocations commonly associated with childhood leukemia; expression of these genes was de-repressed by inhibitors of repressive chromatin modifiers (17). Collectively, these data make it tempting to speculate that the leukemic cells under study were once infected with adenovirus and have acquired a persistent epigenetic signature that has been predicted to be a hallmark of cells transformed by a hit-and-run mechanism (20). Because RUNX1 translocations enhanced viral levels during the acute phase of infection it is possible that other activities of the virus are also enhanced. Additional investigation into how the presence of the ETV6/RUNX1 translocation impacts the adenoviruses ability to modulate cellular genes associated with cancer progression is of great interest. A greater understanding of these changes in patterns of methylation and acetylation, and the genes they affect, may inform the development of therapeutic and diagnostic targets for the treatment or detection of leukemia with a potential viral etiology.

Compelling evidence indicates that most chromosomal translocations found in pediatric leukemias occur before birth. This was suggested by the frequent concordance of clonally identical leukemia in monozygotic twins and confirmed by retrospective analysis of archived neonatal blood samples of children who later developed leukemia (87–89). Most of the common translocations of pediatric leukemia have been identified in cord or neonatal blood samples (90, 91). Although chromosomal translocations are often the initiating event, they are usually not sufficient for full-blown malignancy. A genetic analysis of random cord blood samples found the ETV6/RUNX1 translocation in 1%, a frequency 100-fold greater than the frequency of overt leukemia with that fusion gene in the population at large (37), indicating that 99% of the time these expanded clones fail to develop into leukemia. If indeed a virus is responsible for creating these clones of translocation-containing cells, that virus must be present in at least 1% of otherwise normal newborns. By this criterion adenoviruses alone qualify. Studies looking by PCR for between 6 and 8 viruses in amniotic fluids from abnormal pregnancies found that the vast majority (77% and 84% and 64% and 77%) of virus-positive samples were adenovirus (92–95). In addition, while most studies suggest that the percentage of adenovirus-positive samples is larger among phenotypically abnormal pregnancies, adenovirus DNA is found in the amniotic fluid of 6% (96), 5.4% (93), 2.6% (97), and 5.1% (98) of otherwise normal pregnancies as well. These numbers compare well with our previous detection of adenovirus DNA in 3.7% of cord blood samples from normal pregnancies (5). Our current data reveals that the presence of ETV6/RUNX1 does not inhibit subsequent infections with adenovirus (Fig 6) and, in fact, may enhance the level of virus present during acute infection. Because both adenovirus and the translocation are known to exist contemporaneously *in utero*, it is also possible that the virus epigenetically causes secondary hits in the translocation-positive cells it infects, which further promotes the development of pre-leukemic clones towards leukemia development. Either way, because the translocation does not allow adenovirus to maintain persistent infection in B lymphocytes, one should not expect to find the virus associated with translocation-positive leukemias. Based on our experimental data, however, this outcome does not indicate that adenovirus was never present during the development of disease.

## Conclusion

Both adenovirus and leukemic translocations can be detected in lymphocytes *in utero* and in young children. The results of the current study provide support for how adenovirus could be lost from translocation containing lymphocytes and provide evidence that adenovirus can leave a lasting imprint on cells previously infected in the form of an epigenetic echo. Detection of specific viral epigenetic marks, together with leukemic translocations, may prove useful as an early diagnostic risk factor for disease. Additionally, the use of adenoviruses as vaccines and gene therapy vectors has become increasingly common and we provide evidence of HDAC-mediated repression of adenoviral gene expression. Thus, a full understanding of adenovirus infections in lymphocytes could have a broad impact on the use of adenoviral-based therapies.

## Acknowledgments

We would like to thank Ms. Martine Policard for her technical assistance as an undergraduate student for some of the cellular gene expression experiments.

---

Figure S1. Stable expression of ETV6/RUNX1 or RUNX1/MTG8 does not affect the growth of persistently infected B lymphocytic cells. A persistent infection of Ad5dl309 was established in BJAB cells stably transduced with the empty vector or expression vectors for the indicated leukemic fusion genes. At 28 days post infection, cell cultures were established at 1x10^5^ cells per ml. The number of viable cells was determined daily for 12 days without supplementing the growth medium. The population doubling time (PDT) was calculated by a log-linear regression for the values up to 10 days. Expression of the leukemic fusion transcripts was confirmed in these same cultures and shown in Fig. 1.

