## Supplementary material for "Leukemic fusion genes repress viral gene expression and expel adenovirus from persistently infected human B lymphocytes but evidence of the virus lingers behind": Figure S1

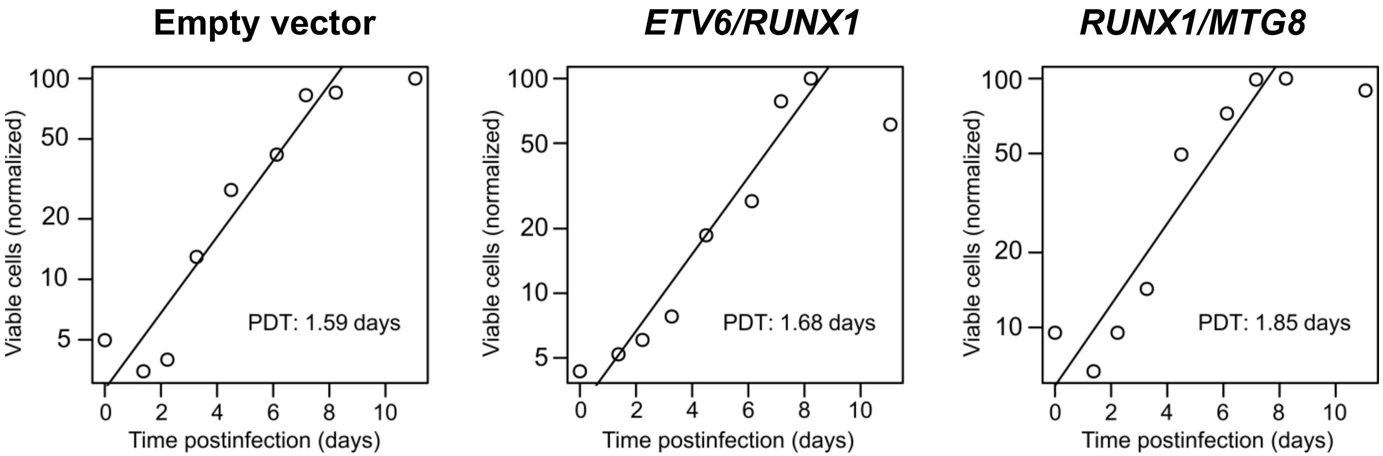


**Figure S1.** Stable expression of ETV6/RUNX1 or RUNX1/MTG8 does not affect the growth of persistently infected B lymphocytic cells.
